# Cell-free expression and biochemical characterization of polysaccharide-synthesizing glycosyltransferases

**DOI:** 10.1101/2024.02.12.580006

**Authors:** Dharanidaran Jayachandran, Amar D. Parvate, Jory T. Brookreson, James E. Evans, Shishir P.S. Chundawat

## Abstract

Polysaccharides are a major class of natural polymers found abundantly across all major life forms and play a critical role as structural, metabolic, or functional components in biomolecular processes. Some polysaccharides like cellulose and hyaluronan are synthesized by membrane-bound family-2 glycosyltransferases (GTs). Despite the fact that the GT-2 family has the maximum number of deposited sequences, the biochemistry of GT-2 family enzymes is still poorly understood due to difficulties associated with GT membrane protein expression, purification, and reconstitution in lipid carriers. Here, we chose *Populus tremula x tremuloides* cellulose synthase 8 (PttCesA8) and *Streptococcus equisimilis* hyaluronan synthase (SeHas) as putative family-2-GTs to be expressed in a wheat-germ-based cell-free expression (CFE) system as proteoliposomes. The cell-free products were obtained as reconstituted liposomes directly from CFE reactions at high yields and short processing times compared to other approaches. GT enzymes expression was confirmed using SDS-PAGE and immunoblotting, and the integration of GTs in lipid layers was observed using cryogenic electron microscopy. Both GTs tested were catalytically active when incubated with their respective substrates and cofactors. The Michalis-Menten kinetic constants, K_m_ for PttCesA8, was 295.8 µM, and SeHas was 321.51 µM (toward UDP N-Acetyl Glucosamine) and 207.88 µM (toward UDP Glucuronic Acid), respectively. UDP was found to actively inhibit both these GTs with apparent inhibition constants of 10.08 µM and 24.38 µM. Mutation of specific conserved residues in structure-deficit SeHas confirmed the importance of lysine-139, glutamine-248, and threonine-283 residues in hyaluronan biosynthesis. In summary, wheat-germ-based CFE can be used to express functionally active and liposome-reconstituted family-2 GTs at high yields with relative ease to enable classical enzymology assays and will also enable more detailed structural studies in the near future.

## 1. Introduction

Glycosyltransferases (GTs) catalyze the transfer of sugar moieties from activated donor molecules to specific saccharide and non-saccharide acceptor molecules, forming glycosidic bonds^1^. According to the Carbohydrate-Active enZYmes (CAZy) database (http://www.cazy.org), there are 116 sequence-based families of GTs that utilize nucleotide diphospho-sugars, nucleotide monophospho-sugars, and sugar phosphates including dolichol-phospho-sugars, sugar-1-phosphates, and lipid diphospho-sugars as activated donors^2, 3^. Among these families, the GT2 family contains the maximum number of sequences originating from bacteria, fungi, plant, and animal species^4^. Some members of family-2 glycosyltransferases include cellulose synthase, hyaluronan synthase, chitin synthase, mannan synthase, mannosyltransferase, glucosyltransferase, galactosyltransferase, and rhamnosyltransferase^1, 5^. The glycosylation reaction in these GTs can proceed with either inversion or retention of stereochemistry at the C1 position of the nucleotide sugar. Some of these GTs combine both polymerizations of nucleotide sugars in an S_N_2-like nucleophilic displacement reaction and translocate the growing polymer across the plasma membrane through its own transmembrane pore^6, 7^. The glycan polymers synthesized by these GTs contain tens of thousands of monomeric sugar units and can be extremely long^4, 8^. Despite being the family with the most significant number of sequences, many of these GTs remain uncharacterized to date. Understanding the structure, mechanism, and function of uncharacterized GTs is crucial in unraveling the mechanism of glycans synthesis and its association with other cellular components.

Cellulose and hyaluronan or hyaluronic acid (HA) are two such polysaccharides synthesized by membrane-bound family-2 GTs and have widespread relevance in bioenergy, biomaterials, and biomedical fields^9,10^. Cellulose is made up of bundles of β-1,4-glucan polymer chains and is the most abundant biopolymer on earth^11, 12^. It is the main structural component of plant cell walls and has a variety of applications in pharmaceutical, food, paper, textile, and furniture industries^4, 13, 14^. In the last few decades, proteins associated with cellulose and hemicellulose synthesis have gained focus with relevance to the production of sustainable biofuels and food. In particular, plant cellulose synthases (CesAs) have been characterized biochemically to a great extent^15–18^. The first structure of homotrimeric plant cellulose synthase was solved by Purushotham et al^19^. Yet, understanding how these CesAs coordinate and work together as a part of cellulose synthase complex (CSC) is still far from fruition both *in vivo* and *in vitro*^20, 21^.

Compared to cellulose, which is a homopolysaccharide, hyaluronan or hyaluronic acid (HA) is an extracellular linear heteropolysaccharide containing alternating N-acetyl-glucosamine (NAG) and glucuronic acid (GA) residues linked via β-1,4 and β-1,3 glycosidic bonds^22^. It is ubiquitously expressed in vertebrates, affecting various physiological processes, including cell adhesion, migration, and differentiation. Recently, Maloney et al^23^ reported the structure of a viral hyaluronan synthase (Has) homologous to vertebrate Has. Although lacking in plants, certain Gram-positive bacteria also produce HA. For instance, Has from *Streptococcus equisimilis* (Se) shares 22-30% sequence similarity with human Has isoforms 1, 2, and 3. Has from bacteria has been characterized extensively using membrane fractions, liposomes, and nanodisc based reconstitution^24–28^. Despite being thoroughly characterized over several decades, the structure of bacterial Has still remains unsolved. However, it has been predicted to have a GT domain, four transmembrane helices, and three amphipathic helices^29^. In addition, biochemical characterization of SeHas has revealed the enzyme to exist as a homodimer, with each monomer responsible for alternating integration of GA and NAG^28^.

One of the major bottlenecks for detailed enzymology and GT structural analysis remains the production of the full-length target membrane proteins with high purity and yield. Previous studies on these GTs have mostly been confined to expression in heterologous expression systems such as bacteria, yeast, and insect cells^15–19, 25, 26, 30, 31^. These studies typically involve the expression, isolation, purification, and reconstitution of GTs into lipid carriers. Heterologously expressed proteins are sometimes toxic to hosts and are also lost during processing steps, often leading to very low yields^15, 32^. To overcome these issues and minimize the total number of processing steps, cell-free protein synthesis technologies have been employed in recent years. Cell-free expression (CFE) systems are robust and customizable expression platforms for producing intact functional proteins with higher yields^33–35^. Recently, cell-free expression systems have been utilized for expressing glycosyltransferases from plants, animals, and bacteria^36–38^.

Here, we used a wheat germ-based cell-free expression system with asolectin liposomes to express and characterize two different membrane-bound family-2 processive GTs – cellulose synthase (EC 2.4.1.12) from *Populus tremula x tremuloides* (Ptt) and hyaluronan synthase (EC 2.4.1.212) from *Streptococcus equisimilis* (Se) that are involved in the synthesis of cellulose and hyaluronan, respectively. Expression of these GTs by CFE was confirmed under immunoblotting using anti-HIS antibodies specific to the proteins. Both these GTs produced uridine diphosphate (UDP) upon incubation with their respective nucleotide sugar substrates (UDP-Glucose for PttCesA8; UDP-GA and UDP-NAG for SeHas) and divalent cations as cofactors (Mn^2+^ for PttCesA8 and Mg^2+^ for SeHas). Using kinetic analyses and product inhibition studies, we determined various reaction parameters specific to the individual GTs. Mutation of certain residues in the SeHas (K139R, Q248A, and T283A) hinders HA biosynthesis and drastically reduces its activity in line with previous report on those mutations^29^. Integration of the GTs into liposomes was confirmed using cryogenic electron microscopy. Our results clearly demonstrate that an established and commercially available wheat germ cell-free expression system could readily express wild-type and mutant GTs belonging to diverse prokaryotes and eukaryotes while maintaining their functional activity, making CFE suitable for readily producing proteoliposomes for detailed GT structural studies in the future directly in proteoliposomes or nanodiscs.

## 2. Materials and methods

### 2.1 Bacterial strain, plasmid constructs, and reagents

pEU-E01 cell-free wheat germ expression vector containing ampicillin resistance and cell-free wheat germ expression kit (Cat# CFS-CPLE-BD Proteoliposome BD Kit) were obtained from CellFree Sciences Co., Ltd. The hyaluronan synthase (*Has*) gene from *Streptococcus equisimilis* carrying a C-terminal 6X-HIS-tag was custom synthesized from Twist Bioscience into the pET28a vector between the NcoI and XhoI restriction sites. Phusion Master Mix was obtained from Thermo Fisher Scientific. Dpn1 and T4 DNA polymerase was obtained from New England Biolabs. Chemically competent *E. cloni* 10G cells were procured from Lucigen. All the primers used in this study were obtained from Integrated DNA Technologies, and DNA sequencing was performed by Azenta Life Sciences (New Jersey). Unless mentioned otherwise, all other reagents were obtained from Sigma Aldrich, VWR, and Thermo Fisher Scientific.

### 2.2 Cellulose Synthase and Hyaluronan Synthase Plasmid Construction

Cellulose synthase 8 (*CesA*8) gene from hybrid aspen (*Populus tremula x tremuloides*) carrying a C-terminal dodeca-HIS-tag from the previously described vector^15^ was moved to the pEU-E01 cell-free wheat germ expression vector using sequence and ligation independent cloning (SLIC) protocol as described previously^21, 39^. Similarly, the hyaluronan synthase (*Has*) gene from *Streptococcus equisimilis* carrying a C-terminal 6X-HIS-tag was moved from the pET28a vector to the pEU-E01 vector. The pEU-E01 plasmid map along for both the genes are shown in Figure S1. Protein sequences for CesA8 and Has are provided in Supplementary Information text S1. The pEU-E01-SeHas mutants (K139R, Q248A, and T283A) were generated using primers designed for site-directed mutagenesis. Sequences of all plasmids (pEU-E01-PttCesA8, pEU-E01-SeHas, pEU-SeHas-K139R, pEU-SeHas-Q248A, pEU-SeHas-T283A) were confirmed using Sanger sequencing method at Azenta (NJ). All the primers used in this study for sequencing, SLIC, and mutations are tabulated in Supplementary Table S1. After sequence confirmation, these plasmids were stored at −80°C for long-term storage. Alternatively, 15% glycerol stocks of *E. cloni* 10G cells carrying the plasmid of interest were maintained at −80°C for future use.

### 2.3 Cell-free production of glycosyltransferases

Cell-free expression plasmids were isolated from *E. cloni* 10G cells using maxiprep kits (Zymo Research, Cat# D4202). Briefly, 150 mL of overnight bacterial culture was used to isolate 100-500 µg of plasmid according to the manufacturer’s specifications. The plasmids were diluted to 1 µg/µl final concentration and were used in cell-free expression reactions. Using the template plasmid, in vitro transcription and translation was completed using the Proteoliposome BD (Bilayer and Dialysis) kit from CellFree Sciences (Product# CFS-CPLE-BD). The transcription reaction was completed at 37°C for 6-hours using a thermocycler. Transcribed mRNA was then used for the translation reaction at room temperature for 72-hours in a bilayer and dialysis arrangement within Slide-A-Lyzer^TM^ MINI Dialysis Devices, 10K MWCO (ThermoFisher Scientific, Catalog#: 88404). Samples were harvested post translation reaction by pipetting and transferred into 1.5 mL micro centrifuge tubes, a small aliquot was saved as a crude sample for gel analysis. Next, 1X TBS (50 mM Tris-HCl, 150 mM NaCl, pH 7.5) was used to rinse the dialysis cup to remove any stuck proteoliposomes before being added to harvested sample. Proteoliposomes were centrifuged at 15,000g for 10-minutes at 4°C to separate the proteoliposome pellet from the CFE supernatant. Supernatants were stored at 4°C for gel analysis. The proteoliposome pellet was washed three times with 1X TBS before being resuspended in 1X TBS to a final volume of 0.5 ml and stored at 4°C. Absorbance at 280 nm measurements of proteoliposomes samples were monitored using Nanodrop2000.

### 2.4 SDS-PAGE gel quantification of cell-free expression products

Protein expression was confirmed by SDS-PAGE analysis using 10% precast gels (Mini-PROTEAN TGX Precast Protein Gels, Bio-Rad, Cat# 4561033). The gels were run for 55-minutes at 150V and stained with G-250 Coomassie stain (Bio-Rad, Cat# 1610803). Final quantified protein yield was determined by gel band intensity measurements using Fiji (https://fiji.sc/).

### 2.5 Immunoblot analysis

Immunoblot analysis was performed, as mentioned previously^15^. Briefly, the cell-free expressed crude and pelleted products were subjected to SDS-polyacrylamide gel electrophoresis. The gel was equilibrated in transfer buffer (25 mM Tris, 190 mM Glycine, and 20% Methanol) for 10 mins before carefully transferring onto a nitrocellulose membrane (Bio-Rad, Cat# 1620112) of dimensions (8.6 X 6.7 cm) using a Bio-Rad Mini-Transfer Cell (Bio-Rad, Cat# 1703930) at 100V with a constant current for 60 mins at 4°C. The transferred nitrocellulose membrane was blocked with blocking buffer (3% (w/v) BSA in PBS-Tween 20 solution) overnight at 4°C. After 16 hours of blocking, the membrane was washed six times (5 mins each) with gentle agitation at 25°C with PBS/Tween 20 buffer. The membrane was later incubated with anti–HIS primary mouse antibodies (1:1000) at 25°C for 1 h. The membrane was washed thrice (5 mins each) with gentle agitation at 25°C with PBS/Tween 20 buffer. After washing, the membrane was incubated with HRP-conjugated anti-mouse secondary antibody (1:1000) and streptactin antibody (specific to the standard protein marker – Bio-Rad, Cat# 1610376; 1:1000 dilution) for 1 h at 25°C before washing it six more times. The membrane was later equilibrated in 1x PBS for 5 mins. Finally, the membrane was incubated with clarity western ECL substrate (Bio-Rad, Cat#1705060) for five more mins before imaging using a chemiluminescence imager.

### 2.6 Cellulose synthase activity assays

Standard cellulose synthase assays were set up as described previously with slight modifications^15^. For the time-course study, twenty microliters of proteoliposomes containing 0.25 µM PttCesA8 was incubated in the presence of 3 mM UDP-Glucose, 20 mM MnCl_2_, in a buffer containing 20 mM Tris (pH 7.5), 100 mM NaCl, and 10% (vol/vol) glycerol. Reactions were incubated at 37°C for 5 h, and samples were collected at regular time points. The samples were centrifuged at 15,000 rpm for 20 mins, and the supernatant was collected and stored for UDP-Glo assay.

Kinetic analyses were performed in the presence of increasing concentrations of UDP-Glucose (0-3 mM), 20 mM MnCl_2_, and 0.5 µM PttCesA8 liposomes. A stock solution of 30 mM UDP-Glucose was diluted to the required concentration for the individual experiments. After incubation for 30 mins at 37°C, the samples were treated as mentioned above.

Product inhibition studies for PttCesA8 were performed in the presence of increasing concentrations of UDP (0-0.5 mM). All reactions contained 0.5 µM PttCesA8 liposomes, 0.3 mM UDP-Glucose, and 20 mM MnCl_2_. Samples were incubated at 37°C for 30 mins before collecting their supernatants for further analysis. The synthesized UDP was quantified relative to the control reactions without substrate.

### 2.7 Hyaluronan synthase activity assays

Hyaluronan synthase assays were set up according to Hubbard et al., with minor modifications^28^. For the time-course study, twenty microliters of proteoliposomes containing 0.3 µM SeHas were incubated in the presence of 5 mM UDP-GA and 5 mM UDP-NAG, 20 mM MgCl_2_, in a buffer containing 40 mM Na_2_HPO_4_ (pH 7.5), 75 mM NaCl, and 5 mM DTT. Reactions were incubated at 30°C for 48 h, and samples were collected at regular time points. The samples were centrifuged at 15,000 rpm for 20 mins, and the supernatant was collected and stored for UDP-Glo assay.

Kinetic analyses for SeHas were performed by keeping the concentration of one substrate constant (2.5 mM UDP-GA or UDP-NAG) and varying the other substrate concentration (0 - 5 mM UDP-GA or UDP-NAG), 20 mM MgCl_2_, 5 mM DTT, and 0.5 µM SeHas liposomes. A stock solution of 50 mM UDP-GA and 25 mM UDP-NAG was diluted to the required concentration for the individual experiments. After incubation for 60 mins at 30°C, the samples were treated as mentioned above.

Product inhibition studies for SeHas were performed according to Tlapak-Simmons et al., with minor modifications^26^. Reactions were set up in the presence of increasing concentrations of UDP (0-0.5 mM). All reactions contained 0.5 µM SeHAS liposomes, 0.3 mM UDP-GA, 0.2 mM UDP-NAG, 20 mM MgCl_2_, and 5 mM DTT. Samples were incubated at 30°C for 60 mins before collecting their supernatants for further analysis. The synthesized UDP was quantified relative to the control reactions in the absence of substrate.

### 2.8 UDP-Glo assay

Ten microliters of supernatant from all the above reactions were incubated with 10 µl of freshly prepared UDP nucleotide detection reagent (Promega, Cat#: V6961) according to the manufacturer’s specifications. The samples were incubated at room temperature for 60 mins in 384-well white luminescent plates (Thermo Fisher Scientific, Cat# 164610), and luminescence was recorded using ‘LUM protocol’ in a Spectramax M5 plate reader (Molecular Devices). All the studies were performed in quadruplicates, and error bars shown are standard deviations from the reported mean value.

### 2.9 Data analysis for obtaining various parameters

Preliminary data analysis was performed using Microsoft Excel^TM^ to obtain the UDP produced and the turnover number (k_cat_). The data from inhibition studies were fitted to the exponential model in Microsoft Excel^TM^ to obtain the K_i_ values. The kinetic data were fitted to the Michaelis-Menten model using the nonlinear curve fitting tool in Origin software to get K_m_ and V_max_ values. Curve fitting was done using the Levenberg−Marquardt algorithm with a tolerance of 1e−9.

### 2.10 Cryo-Electron Microscopy analysis of cell-free reaction products

Approximately 3 µL of proteoliposomes for both CesA8 and SeHas were loaded onto glow discharged Quantifoil grids (300 mesh, R1/2), blotted for 3s and plunge frozen on a Leica EM GP2 to prepare samples for cryo-EM imaging. Grids were loaded onto a 300 keV Titan Krios (Thermo Fisher Scientific) cryogenic electron microscope. Movies were collected in super resolution mode or counting mode at 130,000x magnification using a K3 direct electron detector (Gatan Inc.) and a Bioquantum energy filter (Gatan Inc.) with 20 eV slit width, resulting in a pixel size of 0.3395 or 0.6795 Å respectively. An exposure of 1.65s across 41 frames per movie resulted in total dose of ∼45-50 e-/ Å^2^. All movies were collected at a defocus of −1 to −2 µm using standard EPU software. Motion correction was performed using cryoSPARC V4.0^40^ using standard parameters. Images were binned 4x in Fiji for improving contrast and further visualization^41^.

## 3. Results

### 3.1 Expression of Glycosyltransferases Using Cell-Free System

The pEU-E01 plasmids carrying Cellulose synthase 8 from Poplar (PttCesA8) with C-terminal 12X-HIS tag and hyaluronan synthase from bacteria (SeHas) with C-terminal 6X-HIS tag genes were expressed in wheat germ expression system at EMSL as described previously^33–35^. The overall schematic of the cell-free expression system is shown in Figure 1. Protein expression was confirmed using SDS-PAGE and compared to the control reaction mixture with no plasmid (Figure 2). After spinning down the cell-free expressed proteoliposomes, most of the target protein yield was detected in the proteoliposome pellet (Figure 2, Lane “P” for CesA8 and SeHas) compared to the supernatant. The concentration of PttCesA8 and SeHas were estimated using the band intensity on the SDS-PAGE gel and was found to be 0.33 mg/ml and 0.34 mg/ml. These proteoliposomes were used for all reported biochemical assays.

**Figure 1.**
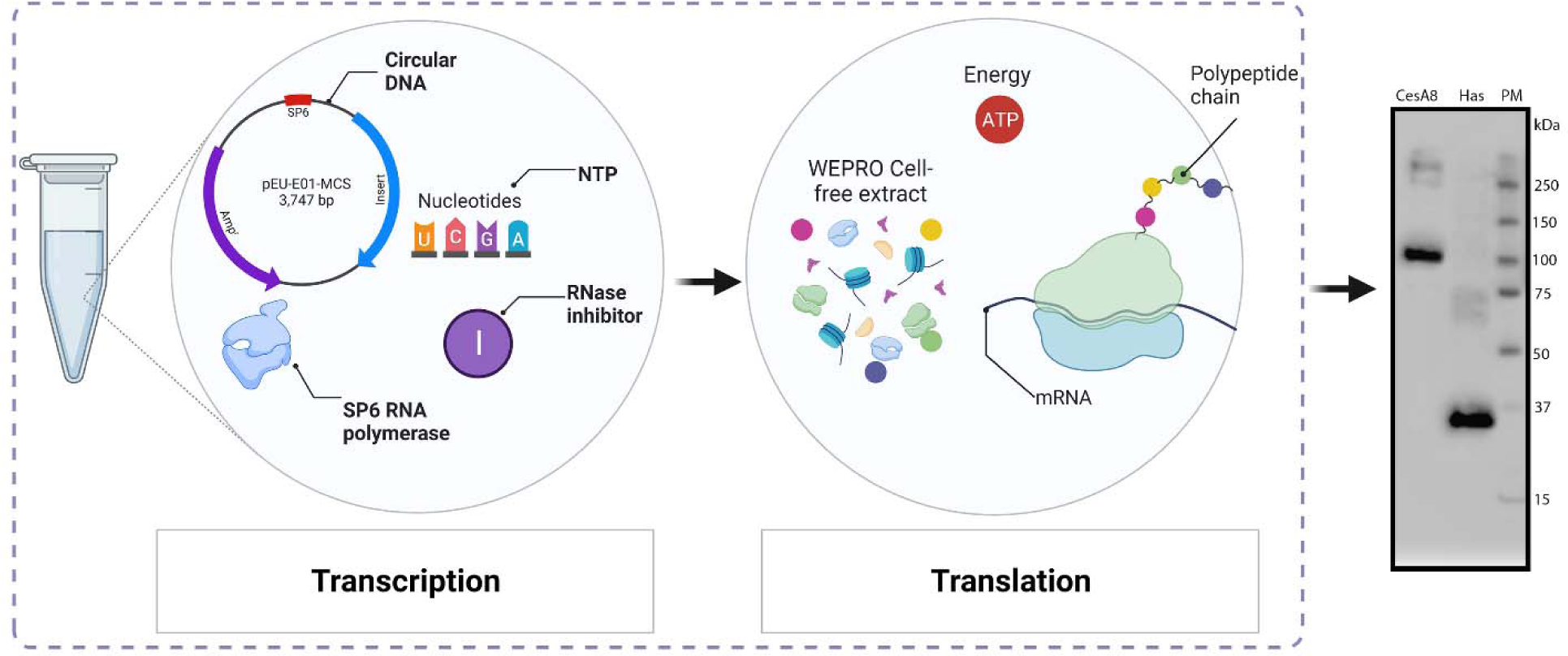
Schematic workflow of cell-free expression method. (a) (*Left*) Transcription reaction in the presence of circular DNA, SP6 RNA polymerase, dNTPs, and RNase inhibitor; (*Middle*) Translation reaction in the presence of WEPRO cell-free extract and creatine kinase; (*Right*) Western Blot-raised against the C-terminal His-tag of CesA8 and HAS. PM - Protein Marker; The image is not to scale.

**Figure 2.**
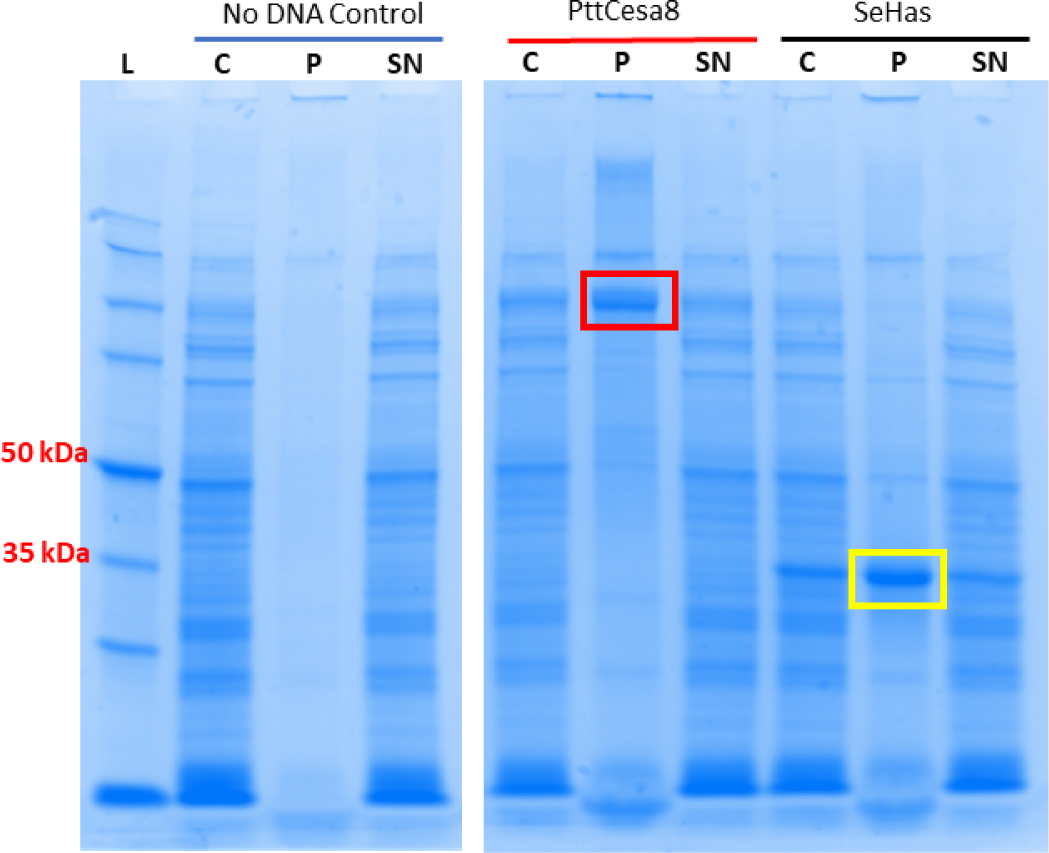
Coomassie blue stained SDS-PAGE of cell-free expressed control reaction lacking DNA and reactions containing DNA encoding PttCesA8 and SeHas proteins. Note the presence of both target proteins in the proteoliposome pellet (red and yellow rectangles) following one-step centrifugal separation and purification. L - ladder, C - crude, P - proteoliposome pellet, SN - supernatant.

The crude reaction mixture and the pelleted liposomes were collected and subjected to immunoblot analysis. As observed under the chemiluminescence imager, the expressed enzymes were immunoreactive to anti-HIS antibodies compared to the control reaction with no plasmids (Figure S2). Blue and red arrowheads represent the position of PttCesA8 and SeHas.

### 3.2 Time Course Study, Kinetic Characterization, and Product Inhibition of PttCesA8

Following previous successes with plant cellulose synthases in *Pichia pastoris* based expression^15^ and characterization, we used reconstituted PttCesA8 liposomes directly from cell-free expression reactions for analyzing the enzyme’s catalytic activity in the presence of UDP-Glucose and Mn^2+^. CesA reactions result in the formation of cellulose alongside UDP nucleotide. The synthesized UDP can be quantified using a luminescence-based UDP glo assay^15, 18^. As shown in Figure 3a, PttCesA8 was active and continued to produce UDP up to 180 mins at 37°C. The maximum amount of UDP accumulated was 380.63 nM. Control reactions with blank liposomes did not produce any UDP, and the background was reduced from the obtained values. The catalytic activity stalls beyond 180 mins, probably due to product inhibition or depletion in the enzyme activity.

**Figure 3.**
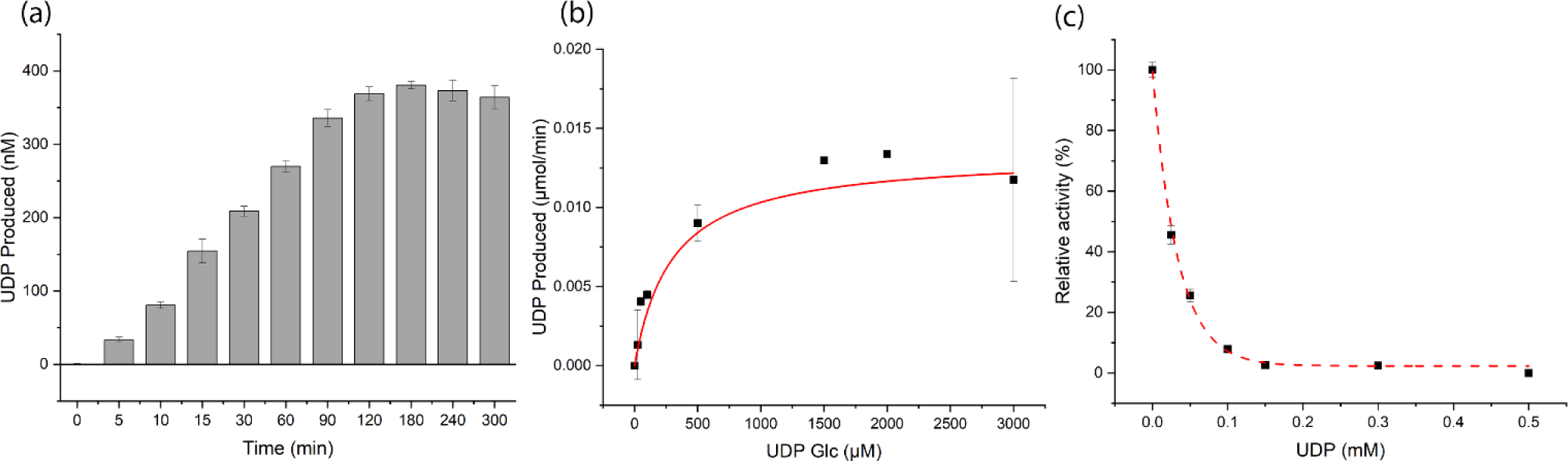
(a) In vitro synthesis of UDP from cell-free expressed CesA8. PLs containing 0.25 μM CesA8 were incubated at 37°C for the indicated time, and the UDP produced was estimated using UDP-Glo assay. (b) Kinetic analysis of cellulose synthesis. Cellulose synthesis reactions were performed with PLs containing 0.5 μM CesA8 in the presence of increasing concentrations of UDP-Glucose (0-3 mM) for 30 min at 37°C. The experimental data (black squares) were fit to monophasic Michaelis-Menten kinetics (red curve). (c) Product inhibition of cellulose synthase. Cellulose synthesis assays were performed with 0.5 μM PL-reconstituted CesA8 in the presence of increasing concentrations of UDP (0-0.5 mM) and 0.3 mM UDP-Glucose for 30 min at 37°C. The obtained product was quantified relative to the UDP formed in the absence of any inhibitor and controls containing no substrate. The experimental data (black squares) was fit to an exponential curve (dashed red curve). All experiments were performed with at least four replicates, and the errors are standard deviations from the mean.

For kinetic analyses, the catalytic activity of PttCesA8 was monitored under increasing concentrations of UDP-Glucose and constant Mn^2+^. PttCesA8 kinetics was observed to follow monophasic Michaelis–Menten kinetics with apparent K_m_ and V_max_ values of 295.8 µM and 0.013 µmol/min, respectively. The experimental data are shown as black squares, and the fitted Michaelis–Menten kinetic curve is shown in red (Figure 3b). The turnover number (k_cat_) was estimated to be 21.57 sec^-1^, and the catalytic efficiency (k_cat_/K_m_) was found to be 0.073 µM^-1^. sec^-1^ (Table 1).

**Table 1.** Summary of kinetic parameters for cell-free expressed PttCesA8 and SeHas. Michaelis-Menten constant (K_m_), maximum velocity (V_max_), turnover number (k_cat_), catalytic efficiency (k_cat_/K_m_), and inhibition constant (K_i_).

| Enzyme<br>(substrate) | $K_m$ ($\mu\text{M}$) | $V_{\max}$ ($\mu\text{mol. min}^{-1}$) | $k_{\text{cat}}$ ($\text{sec}^{-1}$) | $k_{\text{cat}}/K_m$ ($\mu\text{M}^{-1} \cdot \text{sec}^{-1}$) | $K_i$ ($\mu\text{M}$) |
| --- | --- | --- | --- | --- | --- |
| PttCesA8 (UDP-Glc) | 295.8 | 0.013 | 21.57 | 0.073 | 10.08 |
| SeHas (UDP-NAG) | 321.51 | 0.63 | 1042.67 | 3.24 | 24.38 |
| SeHas (UDP-GA) | 207.88 | 0.33 | 546.16 | 2.63 |  |

In order to understand the inhibitory effect of UDP, we examined the catalytic activity of PttCesA8 at increasing concentrations of UDP (0-0.5 mM) and constant UDP-Glucose and Mn^2+^(Figure 3c). The activity of PttCesA8 was estimated relative to the UDP formed in the absence of any inhibitor. In addition, the background arising from UDP added was subtracted using controls with no substrate. The inhibitory constant K_i_ was calculated to be 10.08 µM. A lower value of K_i_ than the K_m_ value suggests that UDP, a reaction product of PttCesA8, is a potent inhibitor and competitively rebinds to the active site. Ultimately, to overcome the UDP inhibition, the substrate (UDP-Glucose) concentration was kept 10 times above the K_m_ value in all reactions.

### 3.3 Time Course Study, Kinetic Characterization, and Product Inhibition of SeHas

Cell-free expressed SeHas proteoliposomes robustly synthesized hyaluronan (HA) when incubated with UDP-GA, UDP-NAG, and Mg^2+^. Similar to CesA, HA catalyzes UDP sugars and forms hyaluronan and UDP as two products. HA synthesis was performed at 30°C until the UDP produced reached saturation. Interestingly, SeHas containing proteoliposomes continued to produce UDP until 24 h, beyond which the product accumulation profile leveled off (Figure 4a). The synthesized UDP was quantified using the UDP Glo assay, as mentioned above.

**Figure 4.**
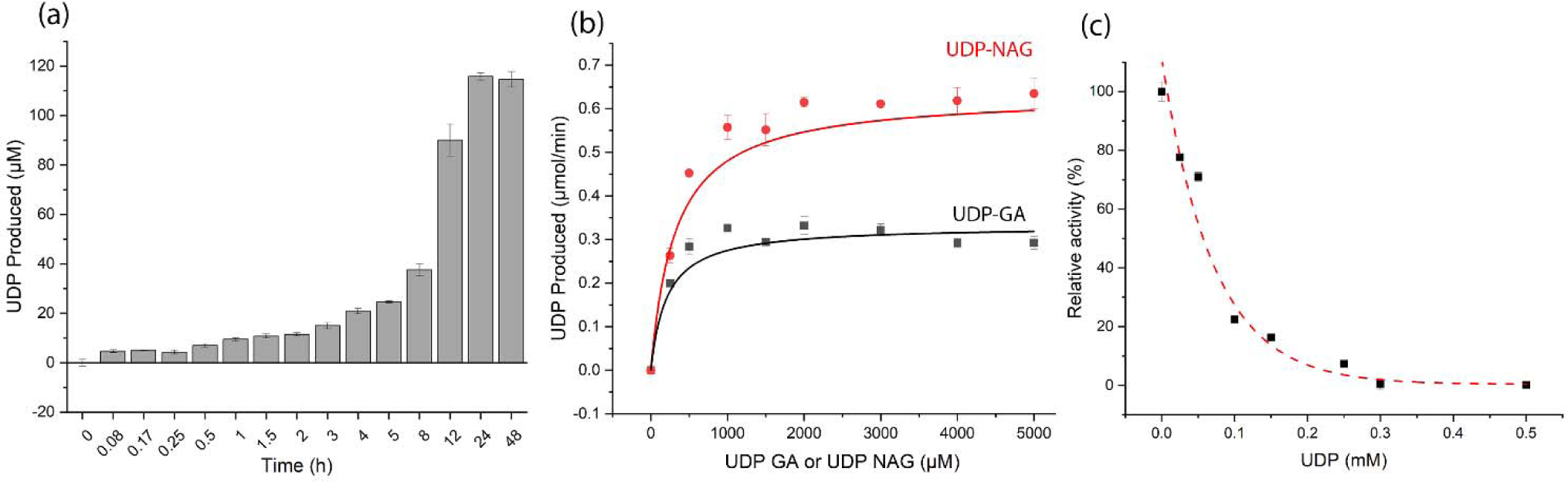
(a) In vitro synthesis of UDP from cell-free expressed SeHAS. PLs containing 0.3 μM SeHAS were incubated at 30°C for the indicated time, and the UDP produced was estimated using UDP-Glo assay. (b) Kinetic analysis of hyaluronan synthesis. Hyaluronan synthesis reactions were performed with PLs containing 0.5 μM SeHAS in the presence of increasing concentrations of UDP-Glucuronic Acid or UDP-N-Acetyl Glucosamine (0-5 mM) while keeping the other substrate concentration constant at 2.5 mM for 60 min at 30°C. The experimental data (black squares) were fit to monophasic Michaelis-Menten kinetics (black curve) for constant UDP-N-Acetyl Glucosamine and varying UDP-Glucuronic Acid. Similarly, experimental data (red circles) were fit to monophasic Michaelis-Menten kinetics (red curve) for constant UDP-Glucuronic Acid and varying UDP-N-Acetyl Glucosamine. (c) Product inhibition of hyaluronan synthase. Hyaluronan synthesis assays were performed with 0.5 μM PL-reconstituted SeHAS in the presence of increasing concentrations of UDP (0-0.5 mM) and 0.3 mM UDP-GA and 0.2 mM UDP-NAG for 60 min at 30°C. The obtained product was quantified relative to the UDP formed in the absence of any inhibitor and controls containing no substrate. The experimental data (black squares) was fit to an exponential curve (dashed red curve). All experiments were performed with at least four replicates, and the errors are standard deviations from the mean.

To obtain the kinetic parameters K_m_ and V_max_, the synthesis of HA was performed by keeping the concentration of one substrate constant (2.5 mM UDP-GA or UDP-NAG) and varying the other substrate concentration (0 - 5 mM UDP-GA or UDP-NAG) and at constant Mn^2+^. Since there are two different substrates, the K_m_ values for each substrate were calculated separately. The experimental data and fitted curve are shown in black for constant UDP-NAG and varying UDP-GA and red for constant UDP-GA and varying UDP-NAG (Figure 4b). K_m_ values for UDP-NAG and UDP-GA were determined to be 321.51 µM and 207.88 µM, respectively. Similarly, the V_max_ values for UDP-NAG and UDP-GA are 0.63 µmol/min and 0.33 µmol/min. The turnover number (k_cat_) was calculated to be 1042.67 sec^-1^ and 546.16 sec^-1^ for UDP-NAG and UDP-GA. The catalytic efficiency (k_cat_/K_m_) for UDP-NAG and UDP-GA was 3.24 µM^-1^. sec^-1^ and 2.63 µM^-1^. sec^-1^ (Table 1).

To assess if UDP also inhibits SeHas activity similar to CesA, we assayed proteoliposomes containing SeHas in the presence of increasing amounts of UDP (0-0.5 mM) and by keeping UDP-NAG and UDP-GA below their K_m_ values, i.e., 0.3 mM UDP-GA and 0.2 mM UDP-NAG. The activity of SeHas reduced significantly with increasing concentrations of UDP with an apparent K_i_ of 24.38 µM (Figure 4c). The activity of SeHas was estimated relative to the UDP formed in the absence of any inhibitor. In addition, the background arising from UDP added was subtracted using controls with no substrate. To ensure the continuous production of HA, the concentration of substrates was kept almost 15-25 times above reported K_m_ values.

### 3.4 Site-directed mutagenesis of conserved residues reduces SeHas catalytic activity

SeHas shares 29.13% and 25.6% sequence identity with Has 2 from *Homo sapiens* (Hs) and Chlorella virus (Cv), respectively (Figure S3). The CryoEM structure of Has from Chlorella virus showed the role of some of these critical conserved residues in priming and substrate binding^23^. Two of these conserved polar amino acids (Lysine-139 and Glutamine-248) were chosen for mutation studies in SeHas. Lysine-139 was mutated to Arginine (K139R), and Glutamine-248 was mutated to Alanine (Q248A) as described elsewhere. We performed the same cell-free expression into proteoliposomes as described earlier (Figure S4) and overall yields were comparable to wild-type. As observed under the UDP-Glo assay, both these mutations led to a complete loss in activity, suggesting a significant functional role of these residues. K139R and Q248A showed only 0.41% and 0.04% activity with respect to the SeHas wild type after 24 h of incubation (Figure 5).

**Figure 5.**
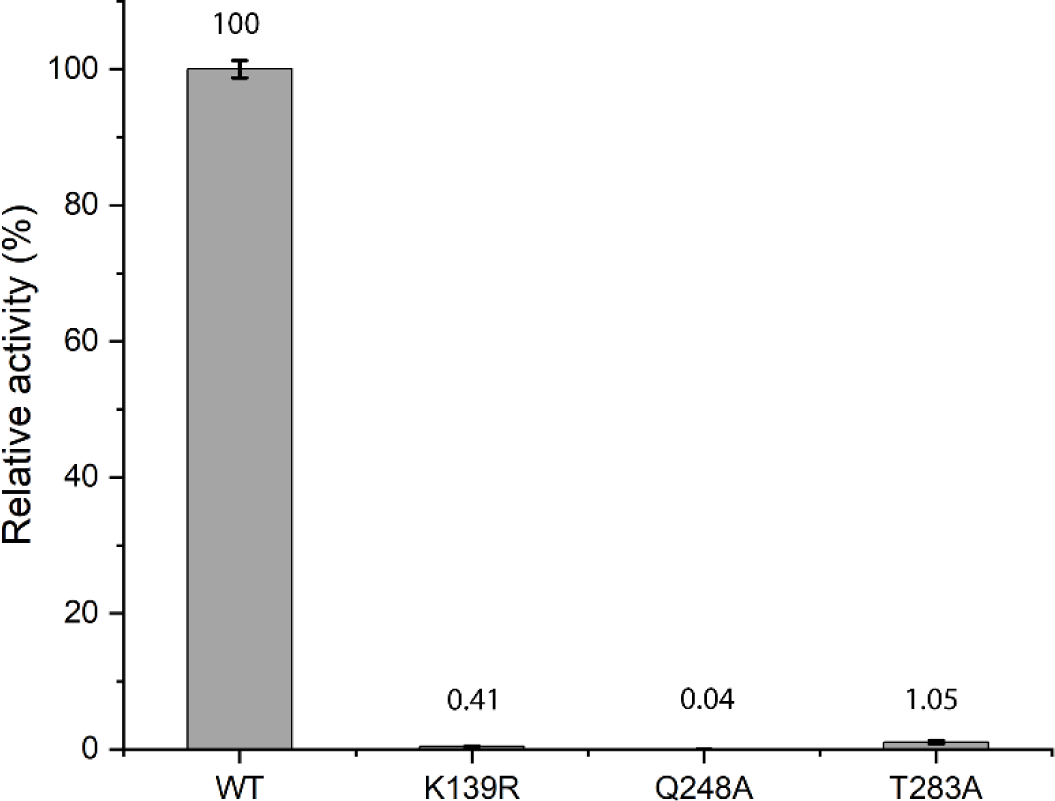
Relative activity (%) of UDP formed in cell-free expressed SeHAS mutants with respect to the wild type. PLs containing 0.3 μM SeHAS were incubated at 30°C for 24 h, and the UDP produced was estimated using UDP-Glo assay.

Similarly, another polar amino acid (Serine-326 in CvHas) is critical in forming a pocket that accommodates the acetamido group of NAG^23^. However, in HsHas and SeHas, this residue is replaced by Threonine, a conserved polar amino acid (marked with ‘*’ in Figure S3). To study if the role of Threonine is similar in bacteria, we mutated Threonine-283 residue to a non-polar amino acid with a simple side chain viz., Alanine. Mutation at this site showed a reduced activity of 1.05% compared to the wild type implying that it could be a critical supporting base residue during HA synthesis.

### 3.5 Visualizing the cell-free products

We employed cryo-electron microscopy to confirm embedding of cell-free expressed GTs in liposomes. A mixture of protein embedded proteoliposomes and empty liposomes of various sizes (20 nm to few hundred nm) were observed in the vitrified samples for both PttCesA8 as well as SeHas samples (Figure S5). Empty liposomes retained the canonical double leaflet lipid bilayer appearance as expected. In comparison, the reactions generated by direct expression in the presence of liposomes resulted in formation of proteoliposomes that exhibited small density protrusions across the lipid bilayer which indicated successful embedding of the GTs. For both proteins, the solvent exposed domains only extended a few nanometers outside the membrane and the exposed domains for SeHas were smaller than for PttCesA8 as anticipated from earlier cryo-EM structures of CesA8 in detergent micelles and CvHas in nanodiscs. In rare instances, dark densities were observed within the lipid bilayer for PttCesA8 (Figure S5b, red boundary) and likely correspond to transmembrane spanning domains. Work is ongoing to solve the structure of these GTs in the lipid bilayer of these proteoliposomes, lipodiscs, or nanodiscs without any exposure to detergents, but that is the focus of a future manuscript exploring structural changes in different environments. The current results clearly demonstrate that the protein is embedded as integral membrane proteins within the lipid bilayer of the asolectin liposomes and the kinetic assays shown above highlight the functional nature of the cell-free expressed GTs.

## 4. Discussion

Cellulose is an important structural cell wall polysaccharide in vascular plants, microbes, algae, and tunicates. This homopolymer is insoluble in water beyond a degree of polymerization of about eight^42^. Owing to its wide range of applications in various industries, it is imperative to understand the synthesis of cellulose at a molecular level. Likewise, hyaluronan is another extracellular matrix polysaccharide found in vertebrates, bacteria, and viruses. It is a heteropolymer that is water soluble, and forms hydrated gels in aqueous solutions. The formation of HA capsules is an effective strategy for pathogens to evade host immune responses^43^.

Cellulose is synthesized by membrane-integrated cellulose synthase complexes (CSC), consisting of an unknown number of individual enzyme monomers (predicted to be a rosette complex of 18-36 individual monomer units in plants)^20^. These enzymes utilize UDP-Glucose as a substrate to synthesize cellulose, elongate the non-reducing end of the polymer, and translocate the synthesized cellulose through the transmembrane pore. Bacterial hyaluronan synthase has been predicted to have a similar mechanism in which the enzyme uses two different substrates: UDP-N-Acetyl Glucosamine and UDP-Glucuronic acid. Recently, the structure of the homotrimeric plant CesA8 from Poplar^19^ and hyaluronan synthase from Chlorella virus^23^ was solved using cryo-electron microscopy, elucidating the mechanisms of family-2 GTs to a great extent. Despite the availability of 3D structures for some GTs, there are still significant gaps in understanding how they coordinate and function at molecular and complex levels.

Cell-free expression systems have been used for expressing drug transporters and other membrane proteins^44, 45^. They offer a significant advantage over the conventional methods of expression and purification since they reduce the processing time of reconstituted membrane protein samples, ultimately leading to higher yield and purity. We performed various biochemical analyses such as time-course, kinetic, and inhibition studies to reaffirm whether the cell-free expressed GTs show similar characteristics to the previously reported studies^15, 28^. PttCesA8 was active for up to 180 mins, beyond which the enzyme’s catalytic activity ceased, consistent with our previous study on plant cellulose synthases^15^. In contrast, it took 90 mins for PttCesA8 and 150 mins for PpCesA5 to reach a plateau in product production in some previously reported studies^16, 17^. Kinetic analysis revealed the K_m_ value for PttCesA8 to be 295.8 µM. Michaelis-Menten constant (K_m_) gives an idea of the affinity of the substrate towards the enzyme. The observed K_m_ is comparable to most previously reported values for cellulose synthases previously, suggesting the cell-free expressed product retains the functional properties of the enzyme^15, 17, 31^. Inhibition constant or K_i_ was observed to be 10.08 µM indicating a strong affinity for inhibitor (UDP) towards the CesA8 active site. A slightly higher K_i_ value (27 µM) has been reported for Poplar CesA8^16^. The turnover number (k_cat_) of PttCesA8 was estimated to be 21.57 sec^-1^, and the catalytic efficiency (k_cat_/K_m_) was found to be 0.073 µM^-1^. sec^-1^ (Table 1). Analogously, SeHas had a prolonged product accumulation (UDP) up to 24 h at 30°C. The enzyme produced almost 115.8 µM of UDP, suggesting a robust turnover mechanism. K_m_ values for UDP-NAG and UDP-GA were calculated to be 321.51 µM and 207.88 µM, respectively, indicating an increased affinity of one substrate over the other. Similar behavior has been observed in HA synthesis where the K_m_ values of UDP-GA are lower than UDP-NAG^24, 26, 27^. HA synthesis was found to be completely inhibited by UDP at a concentration of around 0.3 mM with an apparent K_i_ of 24.38 µM. Nucleotides have been shown to inhibit the activity of SeHas, with 1 mM UDP showing a maximum inhibition of about 97% in one of the previous studies^26^. Additionally, the turnover number (k_cat_) of SeHas was calculated to be 1042.67 sec^-1^ for UDP-NAG and 546.16 sec^-1^ for UDP-GA. The turnover number has mostly been calculated for *Pasteurella multocida* and has been found to be between 7.9 and 13.5 sec^-1^.^46^ Conversely, bacterial hyaluronan synthase has been calculated to have a turnover of 120 sec^-1^ and 150 sec^-1^ for UDP-GA and UDP-NAG, respectively^26^. However, these studies were performed on purified protein extract in detergent micelles and not in stable reconstituted lipid vesicles. Hence, a direct comparison might not be ideal in such cases.

Mutation studies in SeHas show the critical roles of conserved polar residues in Has across different species. Mutations at these sites almost made the enzyme inactive when incubated for 24 h. Glutamine residue in CvHas (Q290) has been found to play a significant role in priming-loop retraction in response to substrate binding. Likewise, lysine (K177 in CvHas) seems to coordinate the substrate binding at the active site^23^. These conserved residues are probably critical in substrate binding and conformation change associated with SeHas during HA synthesis. In one of the previous studies, a complete loss in activity was observed for HsHas when this lysine residue was mutated^47^. Similarly, a significant to complete reduction in HA titer was observed when SeHas carrying these two mutations were expressed in bacteria^29^, implying that the roles of these residues are conserved across species.

Additionally, the exact role of some of the conserved residues was unknown previously. One such residue is threonine, whose function is unknown. Although T283A is not conserved across all species, it is still near the predicted substrate-binding pocket of SeHas^28, 29^. Also, this residue is conserved only across human Has and bacterial Has and is not found in viral Has. CvHas has a serine residue in the place of threonine residue. A complete reduction of SeHas activity upon mutating this residue to a non-polar alanine residue suggests that this residue likely plays a crucial role in the binding of the substrate. Future studies aimed at obtaining high resolution structure of bacterial hyaluronan synthase might elucidate the exact role of these residues and mechanism of hyaluronan formation further.

## 5. Conclusions

Glycosyltransferases (GTs) are a large group of enzymes that are involved in the synthesis of various carbohydrates and carbohydrate-associated molecules^15, 48, 49^. These GTs exist as multimeric complexes and coordinate to synthesize polysaccharides^28, 47, 50–52^. Cellulose and hyaluronan are two such polysaccharides that are synthesized by membrane integrated family-2 glycosyltransferases, namely, cellulose synthase and hyaluronan synthase. Despite being the most prominent family of GTs, structural studies on this family are relatively sparse, mainly due to the difficulties involved in the expression and isolation of full-length GTs. Therefore, more straightforward methods are required to express and isolate GTs at optimal yields with minimal handling and impact on their catalytic activity. In this study, we used the wheat-germ-based cell-free expression method to express GTs from both eukaryotes and prokaryotes and performed various biochemical analyses.

In conclusion, our work confirms that different GTs from different kingdoms involved in the polysaccharide synthesis could be readily expressed and reconstituted in liposomes. The cell-free expressed GTs showed similar biochemical and kinetic characteristics to previous studies. Our reported method directly gives a liposome-reconstituted enzyme without the need to purify and reconstitute them into lipids separately. The developed method has broader applications to speed up research focused on biochemical and structural studies of GTs in general.

## Author Information

### Author Contributions/credit statement

D.J. and S.P.S.C. designed all biochemical experiments; A.D.P., J.T.B., and J.E.E. designed cell-free expression and EM experiments; D.J. and A.D.P. performed all experiments; D.J. prepared the original draft of the manuscript; D.J., A.D.P., J.T.B., J.E.E., and S.P.S.C. contributed to analyzing and writing the manuscript. All authors have given approval to the final version of the manuscript.

## Acknowledgement and Funding Sources

This work was supported by the U.S. Department of Energy (Award Number: DE-SC0019313) and Rutgers University. We are grateful to Prof. Jay Sy for access to the chemiluminescence imager. Cell-free expression and cryo-EM work were performed at the Environmental Molecular Science Laboratory (EMSL), a Department of Energy (DOE) Office of Science User Facility sponsored by the Office of Biological and Environmental Research, under project 60012 (https://doi.org/10.46936/reso.proj.2021.60012/60000391) and was supported by DOE Office of Biological and Environmental Research, Biological Systems Science Division (Grant No. FWP 74915; contract No. grid.436923.9).

## Abbreviations

GT: Glycosyltransferase
CFE: Cell-Free Expression
CSC: Cellulose Synthase Complex
CesA: Cellulose Synthase
Ptt: *Populous tremuloides x tremuloides*
HA: Hyaluronic acid or Hyaluronan
Has: Hyaluronan Synthase
Se: *Streptococcus equisimilis*
Hs: *Homo sapiens*
Cv: Chlorella virus
UDP: Uridine Di-Phosphate
GA: Glucuronic acid
NAG: N-Acetyl Glucosamine
MP: Membrane Protein

