## Supplementary information for "Cell-free expression and biochemical characterization of polysaccharide-synthesizing glycosyltransferases"

**This SI file includes the following:**

Figures S1 to S5 (Pages 2-4)

Supplementary Table S1 (Page 5)

Supplementary Text S1 (Page 6)

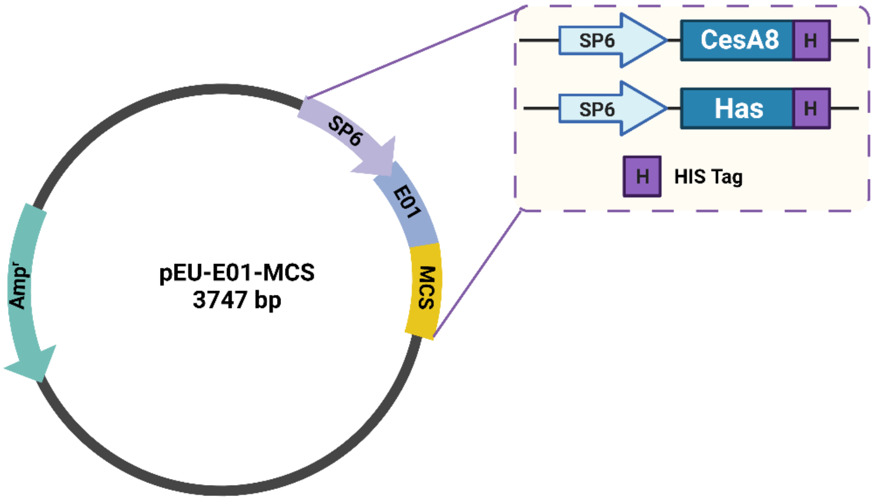


**Figure S1:** Plasmid map for pEU-E01-MCS indicating the insertion of SP6 promoter region, CesA8 followed by 12x-HIS tag, and Has followed by 6x-HIS tag.

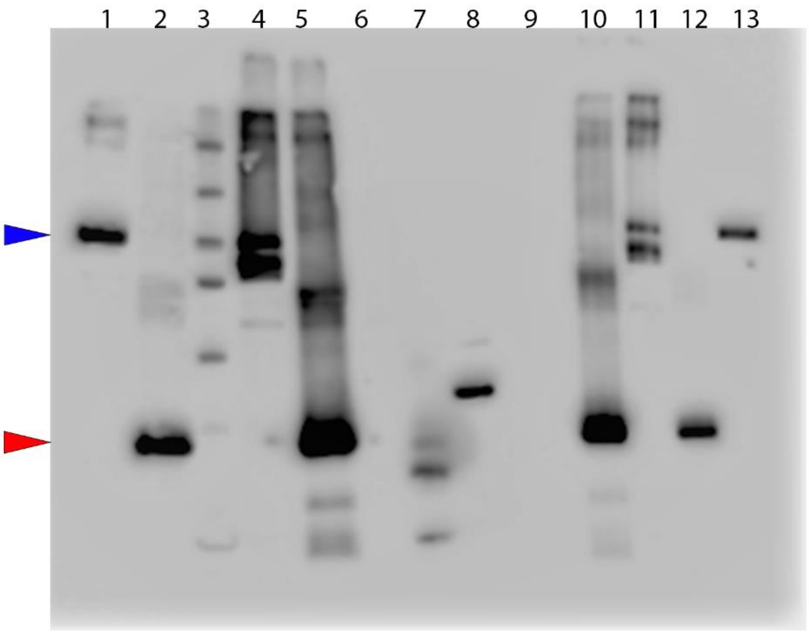


**Figure S2:** Western blot as shown in Figure 1 without cropping. The membrane was raised against the C-terminal His-tag of PttCesA8 and SeHas. Lanes 1-2: PttCesA8 and SeHas pelleted cell-free products (1.5 µg protein loaded); Lane 3: Protein marker; Lanes 4-5: Crude PttCesA8 and SeHas cell-free products (1.5 µg protein loaded); Lane 6: Cell-free product control with no DNA; Lanes 7-8: His-tagged positive controls of different sizes made in-house; Lane 9: Cell-free product control with no DNA; Lanes 10-11: Crude PttCesA8 and SeHas cell-free products (0.75 µg protein loaded); Lanes 12-13: PttCesA8 and SeHas pelleted cell-free products (0.75 µg protein loaded). Blue and red arrowheads correspond to the positions of PttCesA8 and SeHas on the membrane.

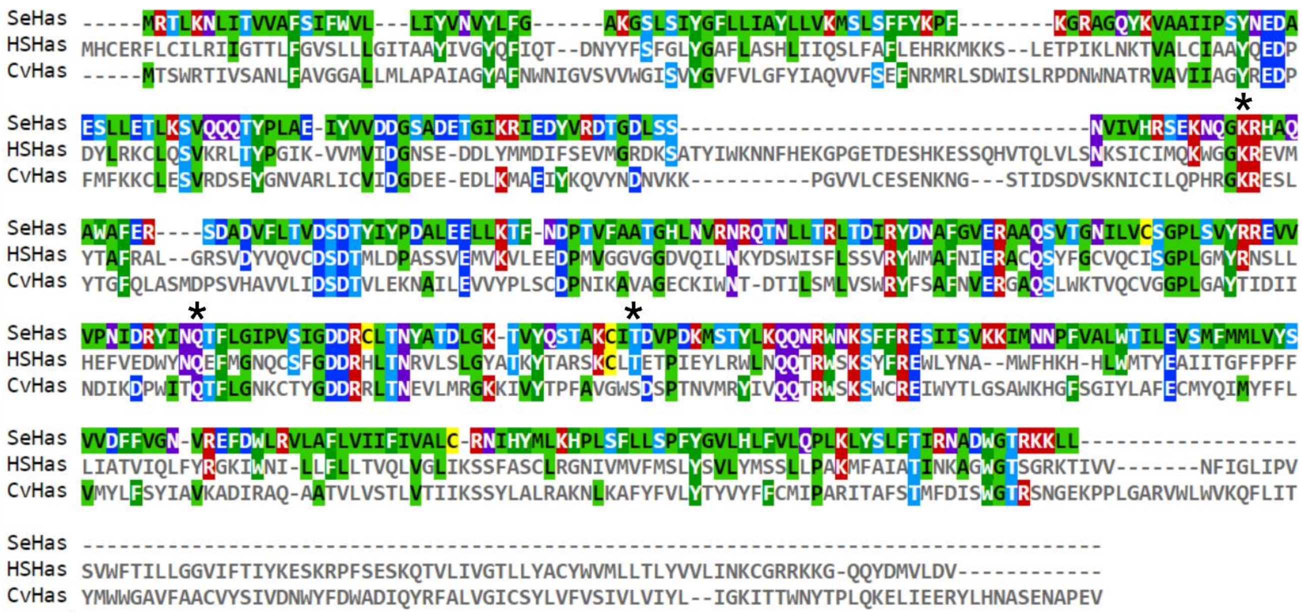


**Figure S3:** Sequence alignment of different hyaluronan synthases. *Streptococcus equisimilis* (Se) Has - GenBank accession no. AAB87874.1, Has 2 from *Homo sapiens* (HS) – NCBI accession no. NP_005319.1, and *Chlorella* virus (Cv) Has - GenBank accession AGE58985.1 were aligned in CLUSTALW and colored based on sequence identity. Mutations (K139R, Q248A, and T283A) in SeHas are marked with an asterisk (*).

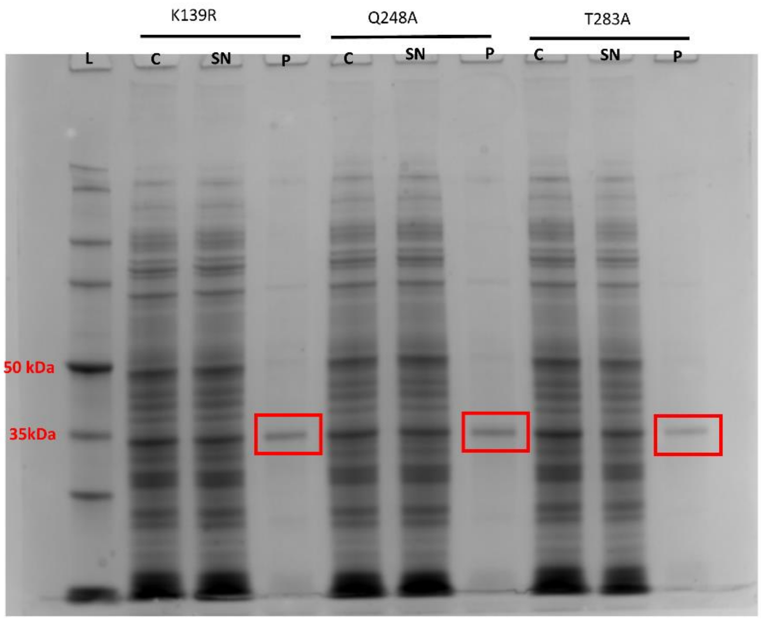


**Figure S4**: Coomassie blue stained greyscale SDS-PAGE of cell-free expressed mutants K139R, Q248A, and T283A. Note the presence of both target proteins in the proteoliposome pellet following one-step centrifugal separation and purification. L- ladder, C – crude, P – Product, SN – supernatant.

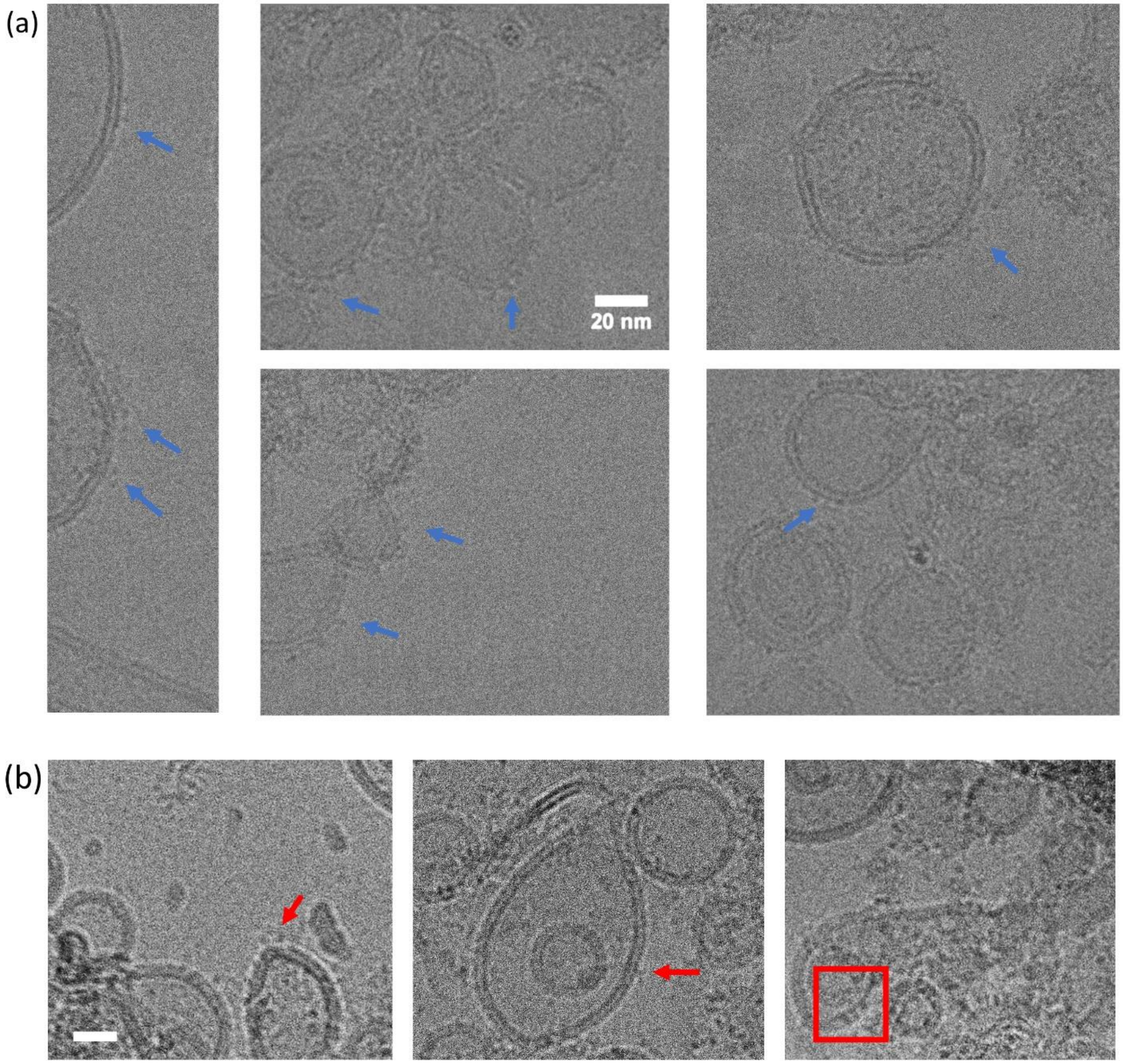


**Figure S5.** CryoEM images of cell-free expressed reaction products showing the integration of transmembrane proteins in liposomes. (a) Blue arrows indicate density attributed to the solvent-exposed domains of SeHas. (b) Red arrows indicate density attributed to solvent-exposed domains of PttCesA8. Scale bar represents 20 nm for all panels.

**Supplementary Table S1**. Primer sequences for target ORF amplification and sequencing.

| Name of Primer | Sequence | Notes |
|---|---|---|
| SP6 promoter, forward primer | GATTTAGGTGACACTATAG | pEU-E01 sequencing primers |
| T7 terminator, reverse primer | GCTAGTTATTGCTCAGCGG | |
| Vec_For_pEU_CesA8 | TCATCATTGAACTAGCCCAAACGAATTCG | SLIC Primers |
| Vec_Rev_pEU_CesA8 | ATTCCATCATTCCCATGGCGATCGCGGC | |
| Ins_For_pEU_CesA8 | CGCCATGGGAATGATGGAATCTGGGGCT | |
| Ins_Rev_pEU_CesA8 | TTGGGCTAGTTCAATGATGATGATGATGAT | |
| Vec_For_pEU_SeHAS | TCACCATTAAACTAGCCCAAACGAATTCG | |
| Vec_Rev_pEU_SeHAS | ATGTTCTCATTCCCATGGCGATCGCGGC | |
| Ins_For_pEU_SeHAS | CGCCATGGGAATGAGAACATTAAAAAAC | |
| Ins_Rev_pEU_SeHAS | TTGGGCTAGTTTAATGGTGATGGTGATG | |
| SeHAS_K139R_For | AAATCAAGGAAGGCGTCATGCAC | Mutation primers |
| SeHAS_K139R_Rev | GTGCATGACGCCTTCCTTGATTT | |
| SeHAS_Q248A_For | GATAGATACATCAACGCGACCTTCCTGGGTATT | |
| SeHAS_Q248A_Rev | AATACCCAGGAAGGTCGCGTTGATGTATCTATC | |
| SeHAS_T283A_For | TAAATGTATTGCAGATGTTCCTG | |
| SeHAS_T283A_Rev | CAGGAACATCTGCAATACATTTA | |

**Supplementary Text S1.** Protein sequences of Poplar Cellulose Synthase 8 and bacterial Hyaluronan Synthase used in this study. All the tags in the proteins are underlined. PttCesA8 has a C-terminal 12x-HIS tag, and SeHas has a C-terminal 6x-HIS tag. Residues chosen for mutations are highlighted in bold red.

>>PttCesA8

MMESGAPICHTCGEQVGHDANGDLFVACHECNYHICKSCFEYEIKEGRKVCLRCGSPY
DENLLDDVEKKGSGNQSTMASHLNNSQDVGIHARHISSVSTVDSEMNDEYGNPIWKNR
VESWKDKRNKKKKSNTKPETEPAQVPPEQQMENKPSAEASEPLSIVYPIPRNKLTPYRA
VIIMRLIILGLFFHYRITNPVDSAFGLWLTSVICEIWFAFSWVLDQFPKWKPVNRETFIERL
SARYEREGEPSQLAAVDFFVSTVDPLKEPPLITANTVLSILAVDYPVDKVSCYVSDDGAA
MLTFESLVETAEFARKWVPFCKKFSIEPRAPEFYFSQKIDYLKDKVQPSFVKERRAMKR
DYEEYKVRVNALVAKAQKTPDEGWTMQDGTPWPGNNTRDHPGMIQVFLGNTGARDIE
GNELPRLVYVSREKRPGYQHHKKAGAENALVRVSAVLTNAPYILNLDCDHYVNNSKA
VREAMCILMDPQVGRDVCYVQFPQRFDGIDRSDRYANRNIVFFDVNMKGLDGIQGPMY
VGTGCVFNRQALYGYGPPSMPRLRKGKESSSCFSCCCPTKKKPAQDPAEVYRDAKRED
LNAAIFNLTEIDNYDDYERSMLISQLSFEKTFGLSPVFIESTLMENGGVPESANSSTLIKEA
IHVIGCGFEEKTEWGKEIGWIYGSVTEDILSGFKMHCRGWRSIYCMPVRPAFKGSAPINL
SDRLHQVLRWALGSVEIFFSRHCPFWYGYGGGRLKWLQRLAYINTIVYPFTSLPLIAYCT
IPAVCLLTGKFIIPTLSNLASMLFLGLFISIIVTAVLELRWSGVSIEDLWRNEQFWVIGGVS
AHLFAVFQGFLKMLAGIDTNFTVTAKAADDTEFGELYMVKWTTLLIPPTTLLIINIVGVV
AGFSDALNKGYEAWGPLFGKVFFAFWVILHLYPFLKGLMGRQNRTPTIVVLWSVLLTS
VFSLVWVKINPFVNKVDNTLAGETCISIDCHHHHHHHHHHHH

>>SeHas

MRTLKNLITVVAFSIFWVLLIYVNVYLFGAKGSLSIYGFLLIAYLLVKMSLSFFYKPFKGR
AGQYKVAAIIPSYNEDAESLLETLKSVQQQTYPLAEIYVVDDGSADETGIKRIEDYVRDT
GDLSSNVIVHRSEKNQG**K**RHAQAWAFERSDADVFLTVDSDTYIYPDALEELLKTFNDPT
VFAATGHLNVRNRQTNLLTRLTDIRYDNAFGVERAAQSVTGNILVCSGPLSVYRREVVV
PNIDRYIN**Q**TFLGIPVSIGDDRCLTNYATDLGKTVYQSTAKCI**T**DVPDKMSTYLKQQNR
WNKSFFRESIISVKKIMNNPFVALWTILEVSMFMMLVYSVVDFFVGNVREFDWLRVLAF
LVIIFIVALCRNIHYMLKHPLSFLLSPFYGVLHLFVLQPLKLYSLFTIRNADWGTRKKLLH
HHHHH
